# Liver organoid-mediated cyclophosphamide neurotoxicity in CNS organoids in a multi-organ microphysiological system

**DOI:** 10.64898/2026.05.17.725752

**Authors:** T. Mitchell, T. Aihara, K. Tanimoto, E. Wolvetang

## Abstract

Cyclophosphamide (CP) is a widely used alkylating agent whose cytotoxic activity depends on hepatic CYP450-mediated bioactivation. While CP-associated neurotoxicity and cognitive impairment are recognized clinically, the mechanisms of secondary organ damage through metabolic cross-talk remain poorly understood due to limitations of conventional monoculture models. Here we employ a multi-organ microphysiological system (MPS) connecting stem cell derived liver and CNS organoids via microfluidic channels to model inter-organ drug metabolism and secondary toxicity. Liver organoids were treated with CP (0-200 µM) for 48 hours, and connected CNS organoids were assessed for secondary damage by confocal Z-stack imaging of DNA damage (γH2AX), neuronal identity (NeuN), and nuclear content (DAPI). We observe dose-dependent reduction in NeuN expression and γH2AX signal in connected CNS organoids, consistent with neurotoxic metabolite transfer from liver. Critically, CNS-to-CNS control connections show no comparable damage at equivalent CP concentrations, confirming that hepatic metabolism is required for CNS toxicity. These findings validate the MPS platform for modelling multi-organ drug toxicity and provide direct evidence that liver-derived CP metabolites drive secondary neurotoxicity through inter-organ metabolic communication.

## Introduction

Cyclophosphamide (CP) is an oxazaphosphorine alkylating agent used extensively in oncology and autoimmune disease management. As a prodrug, CP requires hepatic bioactivation primarily through CYP2B6, with contributions from CYP3A4, CYP2C9, and CYP2A6, to generate the active metabolite 4-hydroxycyclophosphamide, which spontaneously converts to phosphoramide mustard, the ultimate DNA cross-linking and alkylating species^1,2^. This hepatic metabolism also generates chloroacetaldehyde via CYP3A4-mediated N-dechloroethylation, a metabolite implicated in nephrotoxicity and neurotoxicity^3^.

Post-chemotherapy cognitive impairment (PCCI), colloquially termed “chemobrain,” affects a significant proportion of patients receiving CP-containing regimens^4^. Preclinical studies have identified oxidative stress, neuroinflammation, apoptosis, and disruption of neurotrophic signalling as molecular drivers of CP-induced neurotoxicity^3^. However, these studies have predominantly used animal models or monoculture systems that fail to recapitulate the critical inter-organ metabolic communication required for CP bioactivation, since CNS cells alone lack the CYP450 machinery to metabolize CP to its active form.

Microphysiological systems (MPS), also termed organs-on-chips, have emerged as physiologically relevant platforms capable of recreating multi-organ interactions by connecting tissue compartments through microfluidic channels^5,6^. These systems enable the study of metabolic cross-talk between organs, recapitulating scenarios where drug metabolism in one organ generates toxic metabolites that damage another. Recent advances have demonstrated the predictive value of liver-on-chip systems for drug-induced liver injury^7^, while multi-organ configurations have been used to model systemic drug pharmacokinetics^8^.

Human iPSC-derived organoids provide a scalable, human-relevant source of complex three-dimensional tissue models. iPSC-derived brain organoids have been proposed as a model system for chemotherapy-induced CNS toxicity^9^, while liver organoids expressing functional CYP450 enzymes can metabolize prodrugs to their active forms. Combining these organoid systems within an MPS platform creates a unique opportunity to model the complete pathway from hepatic drug metabolism to secondary organ toxicity.

Here we present a multi-organ MPS model connecting stem cell derived liver organoids to CNS organoids via the BioStellar™ Plate, a stirrer-pump-integrated Multi-Organ MPS platform (Sumitomo Bakelite Co., Ltd.). We demonstrate that CP treatment of liver organoids drives dose-dependent DNA damage and loss of neuronal identity in connected CNS organoids, and validate that this effect requires hepatic metabolism through organ-matched controls.

## Results

### Stem cell derived organoids express tissue-specific markers

Liver and CNS organoids were differentiated from a proprietary stem cell line using established protocols (see Methods) and cultured for 90 days prior to MPS assembly. Immunofluorescence characterization of vehicle-treated organoids confirmed robust tissue-specific marker expression. CNS organoids displayed strong NeuN signal (mean intensity 1.5-fold higher than DAPI intensity), indicating mature neuronal differentiation, while liver organoids showed consistent albumin expression (mean 0.8-fold DAPI intensity) across the organoid cross-section (Fig. 1, Fig. 2a). Both organoid types displayed consistent morphology across biological replicates (n=3 per condition), with well-defined spheroid architecture visible in DAPI staining and bright-field imaging. Four wells were excluded from analysis due to absence of organoid signal (see Methods).

**Figure 1.**
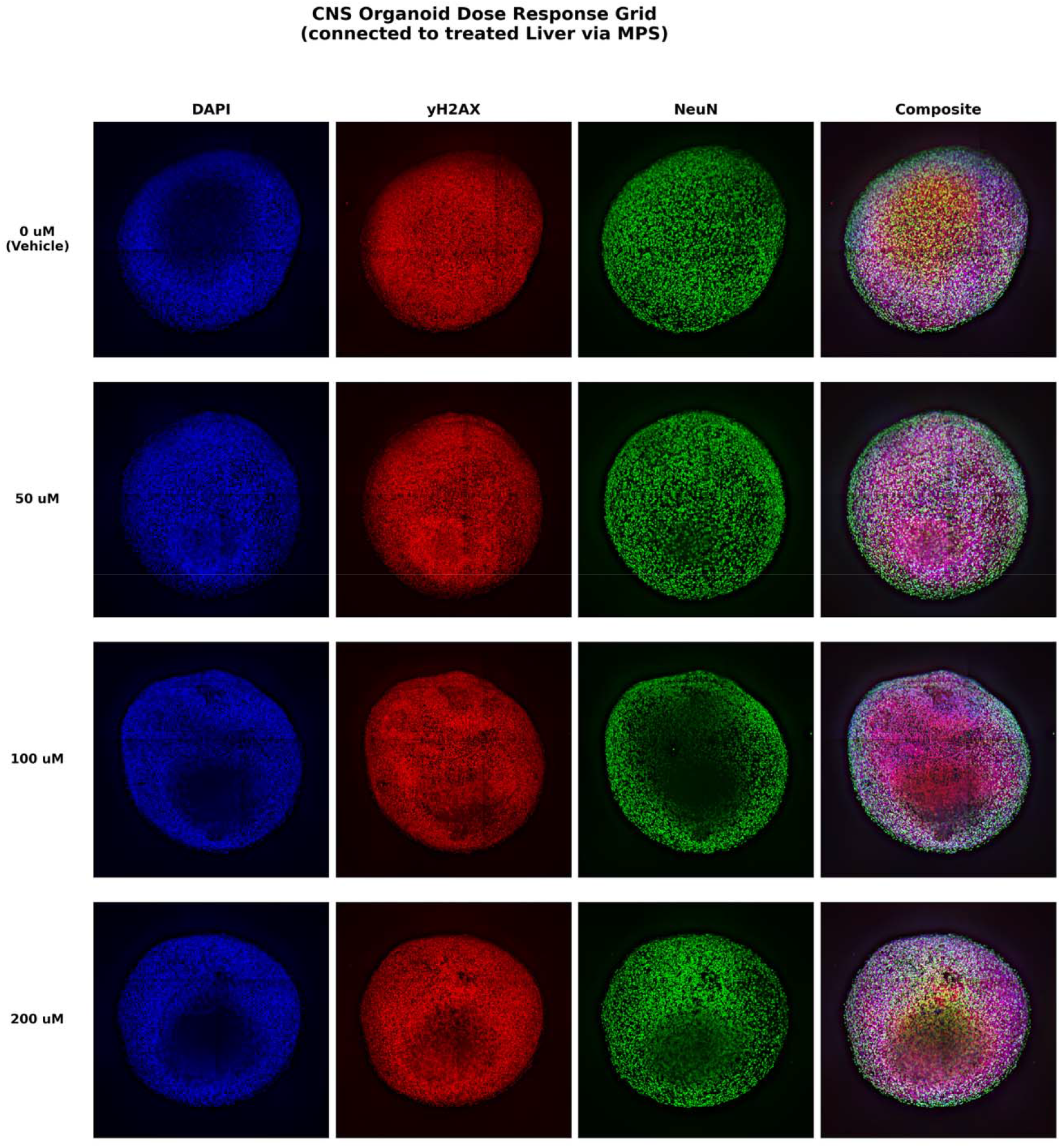
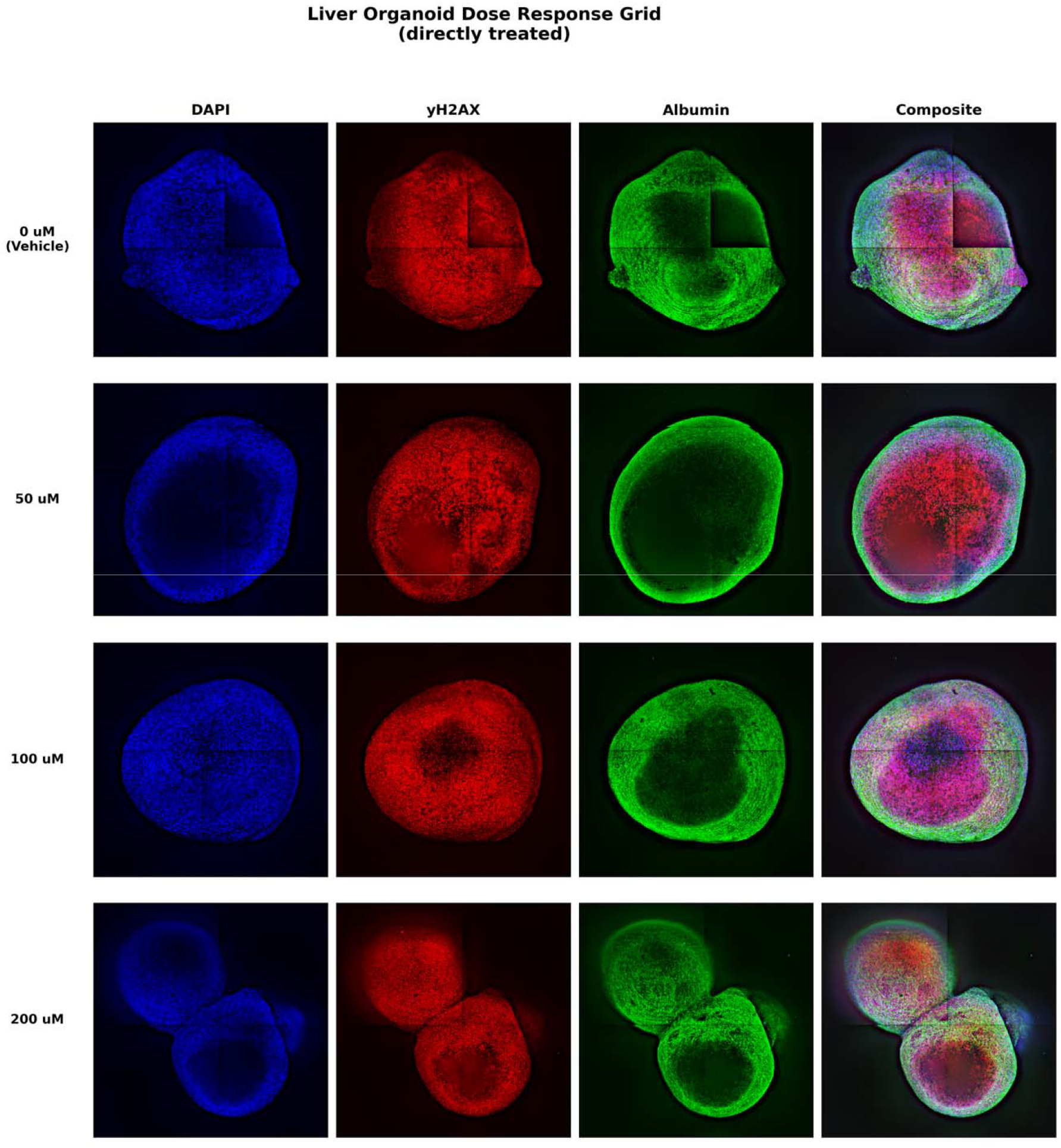
Representative confocal images of CNS and liver organoids across cyclophosphamide dose-response. (Top) CNS organoid dose-response grid showing DAPI (nuclei, blue), γH2AX (DNA damage, red), NeuN (neuronal marker, green), and composite overlay for 0, 50, 100, and 200 µM CP. CNS organoids were not directly treated but connected to treated liver organoids via MPS microfluidic channels. (Bottom) Liver organoid dose-response grid showing DAPI, γH2AX, Albumin (hepatocyte marker, green), and composite for directly treated liver organoids. Images are Extended Depth-of-Field composites generated from ∼40 Z-slice confocal stacks (Yokogawa CQ1, 10x). Representative wells selected by automated quality scoring. Scale: each panel approximately 2 mm across.

**Figure 2.**
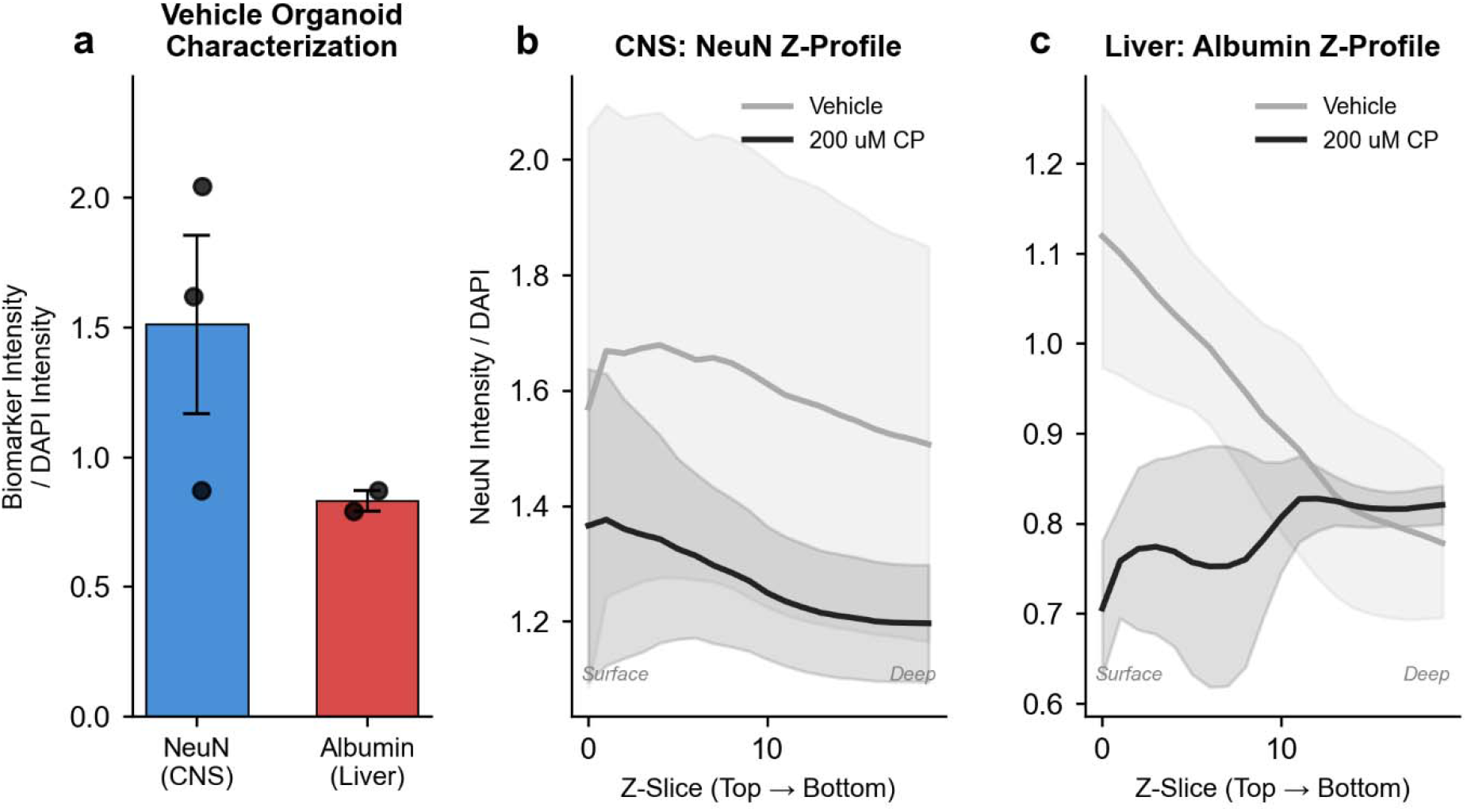
Organoid characterization and volumetric Z-depth profiling. (a) Biomarker intensity normalized to DAPI intensity in vehicle-treated organoids, confirming robust NeuN expression in CNS organoids and albumin expression in liver organoids. Note that absolute values reflect channel-specific imaging parameters (antibody affinity, fluorophore brightness, detector settings) and should not be compared across markers; this metric is designed for within-marker comparison across treatment conditions. Bars represent mean ± SEM; individual data points shown (n=2–3). (b) NeuN intensity per DAPI across the confocal Z-stack (surface to deep) of CNS organoids, comparing vehicle (grey) to 200 µM CP (black). CP-exposed CNS organoids (connected to treated liver) show consistently reduced NeuN signal across the full acquired depth. (c) Albumin intensity per DAPI across the Z-stack of liver organoids. At 200 µM CP, albumin signal diverges from vehicle in the deeper slices, suggesting depth-dependent metabolite penetration and sustained damage in the organoid interior. Shaded regions represent ± SEM. Note: the Z-stack did not capture the full organoid depth for most wells; profiles represent the accessible imaging volume from the organoid surface into the interior.

### Volumetric Z-profiling reveals depth-dependent biomarker reduction

Confocal Z-stack acquisition (∼40 slices per well, every 2nd slice analysed) enabled depth-resolved quantification of biomarker expression from the organoid surface into the interior. Note that the Z-stack did not capture the full organoid diameter for most wells; the profiles represent the accessible imaging volume rather than a complete shell-to-shell transect.

At 200 µM CP, connected CNS organoids showed consistently reduced NeuN intensity per DAPI compared to vehicle across the entire acquired Z-depth (Fig. 2b). This uniform reduction suggests that liver-derived CP metabolites affect neuronal marker expression throughout the organoid rather than being limited to the surface. Liver organoids treated directly with 200 µM CP showed depth-dependent albumin reduction, with signal diverging most prominently from vehicle in the deeper slices (Fig. 2c). This pattern is consistent with metabolite penetration from the surface, with the organoid interior, further from media exchange, showing more sustained effects. These depth-resolved profiles provide spatial information that would be lost in conventional maximum intensity projection analysis.

### Dose-dependent loss of neuronal identity in connected CNS organoids

The primary dose-response experiment examined CP concentrations of 0, 50, 100, and 200 µM applied directly to liver organoids, with connected CNS organoids assessed for secondary damage. The most striking finding was dose-dependent reduction in biomarker intensity normalized to DAPI signal in CNS organoids. NeuN intensity per DAPI decreased from a mean of 1.52 in vehicle controls to 1.14 at 50 µM and 1.19 at 200 µM, representing a 20–25% reduction in neuronal marker expression (Fig. 3a). The slight recovery from 1.14 at 50 µM to 1.19 at 200 µM likely reflects differential cell population dynamics at higher doses: at 200 µM, accelerated apoptotic clearance of the most damaged (lowest-NeuN) neurons may enrich for surviving cells that retain relatively higher NeuN expression, producing a modest rebound in the population-level metric. This non-monotonic pattern is consistent with competing processes of dose-dependent toxicity and dose-dependent clearance operating simultaneously at the 48-hour timepoint. This reduction was observable from the lowest tested dose (50 µM) and persisted through 200 µM. Liver organoids showed a more modest decline in albumin intensity per DAPI from 0.82 (vehicle) to 0.62 (100 µM), though this did not reach statistical significance with n=2– 3 per group.

**Figure 3.**
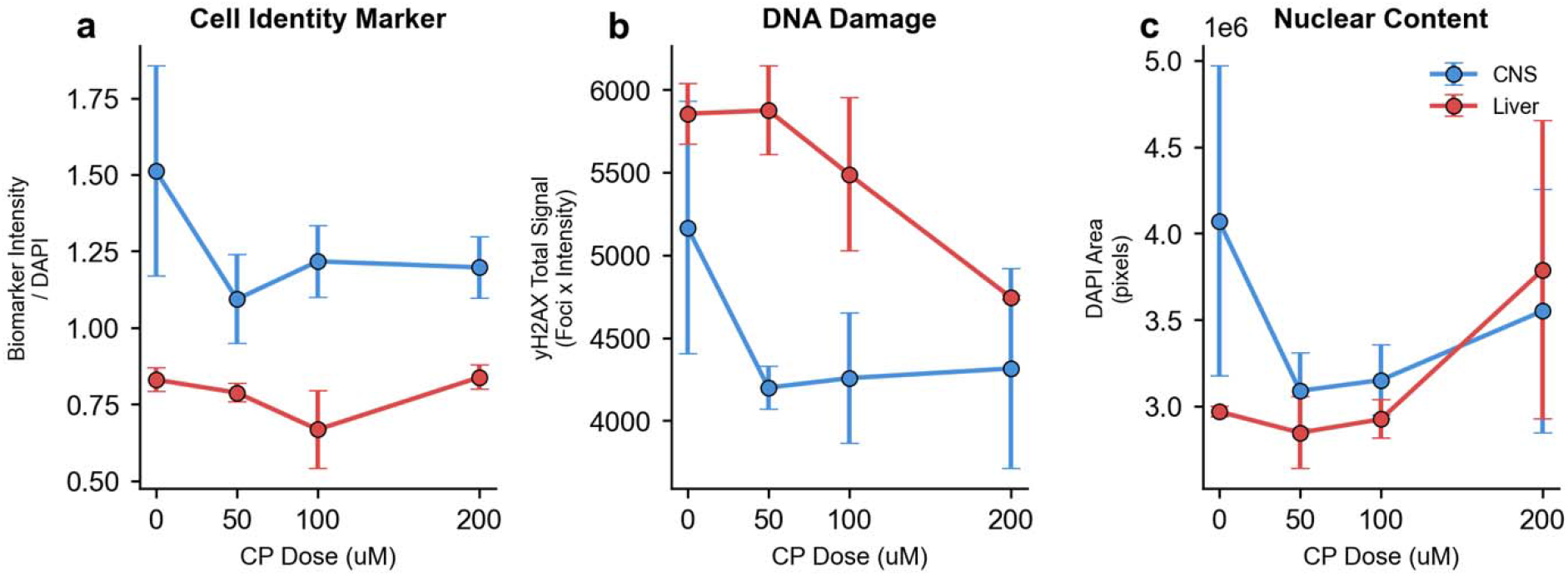
Dose-dependent effects of cyclophosphamide on connected organoids. (a) Biomarker intensity normalized to DAPI signal at the organoid core. CNS organoids (blue) show reduction in NeuN expression with increasing dose of CP applied to connected liver organoids. Liver organoids (red) show modest albumin reduction. (b) γH2AX total signal (puncta density × mean intensity) decreases with dose, consistent with apoptotic clearance of heavily damaged cells by 48 hours. (c) DAPI area at the organoid core as a proxy for nuclear content. Data represent mean ± SEM of core Z-slices (n=2–3 biological replicates per condition).

### γH2AX signal reflects DNA damage and subsequent cell loss

We quantified DNA damage using a composite metric combining γH2AX puncta density (detected via Difference-of-Gaussian blob detection) and mean γH2AX intensity within DAPI-masked regions, providing sensitivity to both discrete foci and diffuse pan-nuclear staining. In CNS organoids, γH2AX total signal decreased from vehicle (mean 5280) to 200 µM (mean 4340), a 17.8% reduction (Fig. 3b). Rather than the expected increase in DNA damage markers with dose, this decrease likely reflects loss of the most damaged cell populations through apoptosis by the 48-hour timepoint, a phenomenon previously described in chemotherapy-treated tissues where initial DNA damage is followed by clearance of heavily damaged cells^10^. Consistent with this interpretation, DAPI area (a proxy for nuclear content and cell viability) showed variable but not statistically significant changes across doses (Fig. 3c), suggesting that while some cell loss occurs, surviving cells exhibit less damage signal per unit area.

Liver organoids showed a similar pattern, with γH2AX total signal declining from approximately 5900 (vehicle) to 4500 (200 µM), consistent with direct metabolite exposure and subsequent elimination of the most damaged hepatocytes (Fig. 3b).

## Discussion

We demonstrate that a multi-organ MPS platform connecting stem cell derived liver and CNS organoids can model liver-mediated secondary neurotoxicity of cyclophosphamide. The key findings are: (1) dose-dependent reduction in NeuN expression in CNS organoids connected to CP-treated liver, indicating loss of neuronal identity; (2) decreased γH2AX signal at higher doses consistent with elimination of damaged cells by 48 hours; and (3) depth-resolved Z-profiling revealing that drug effects penetrate from the organoid surface into the interior, information that would be lost in conventional projection-based analysis.

The reduction in NeuN intensity per DAPI observed in connected CNS organoids aligns with known mechanisms of CP-induced neurotoxicity, including oxidative stress-mediated neuronal damage and disruption of neurotrophic signalling^3^. NeuN (RBFOX3) expression is associated with neuronal terminal differentiation^11^, and its reduction under toxicant exposure may reflect either direct neuronal damage, dedifferentiation, or selective loss of mature neurons. The observation that this effect was detectable from the lowest tested dose (50 µM) suggests sensitivity of the MPS platform to physiologically relevant metabolite concentrations.

The decrease in γH2AX total signal with increasing dose, contrary to the expected increase in a static DNA damage assay, is an important observation warranting discussion. At 48 hours post-treatment, heavily damaged cells have likely undergone apoptosis and been cleared or fragmented, reducing the pool of measurable γH2AX-positive nuclei. This temporal dynamic is consistent with the established progression from acute DNA damage (peak γH2AX at 1–6 hours) through DNA repair or apoptotic commitment (6–24 hours) to cell clearance (24–48 hours)^10^. Future experiments incorporating earlier timepoints (6, 12, 24 hours) and apoptosis markers (cleaved caspase-3) would resolve this temporal sequence.

The organ-matched controls included in this study provide supporting evidence for the MPS platform’s utility. At 50 µM, the Liver-Liver configuration showed γH2AX signal comparable to the main Liver-CNS configuration, consistent with metabolite transfer between connected compartments. The CNS-CNS configuration showed a trend toward reduced damage signal compared to CNS organoids connected to treated liver, consistent with the absence of CYP450-mediated CP bioactivation in neural tissue, although the differences were modest at the 48-hour timepoint, potentially reflecting the same clearance dynamics observed in the dose-response. Future studies with earlier timepoints and larger sample sizes would be needed to fully resolve the kinetics of metabolite transfer between compartments.

Several limitations should be acknowledged. The small sample size (n=2–3 per group after exclusion of empty wells) limits statistical power; future studies should increase biological replicates. The 10x imaging resolution, while sufficient for tissue-level quantification, does not permit single-cell resolution of γH2AX foci, higher magnification imaging would enable per-nucleus foci counting. The confocal Z-stack did not capture the full organoid diameter for most wells, limiting depth profiling to approximately half the organoid volume; future acquisitions should extend the Z-range to encompass the complete organoid. Additionally, the 48-hour single timepoint captures a snapshot of a dynamic process; longitudinal imaging or multiple timepoints would provide a more complete picture of the damage-repair-death kinetics. Conditioned media was collected from treated organoids at the 48-hour endpoint; future analysis of CP metabolites (4-hydroxycyclophosphamide, phosphoramide mustard, chloroacetaldehyde) in these samples by mass spectrometry or functional liver assays would provide direct evidence of metabolite transfer between compartments and strengthen the mechanistic interpretation.

In conclusion, this study provides proof-of-concept that stem cell derived organoids within a multi-organ MPS platform can recapitulate inter-organ drug toxicity pathways relevant to human pharmacology. The demonstration that liver-metabolized CP drives secondary neurotoxicity in connected CNS organoids, validated by organ-matched controls, establishes this platform as a tool for investigating metabolite-mediated multi-organ toxicity in drug development and safety assessment.

## Methods

### Cell culture and organoid differentiation

Liver and CNS organoids were derived from a proprietary stem cell line (The Organoid Company BV) using differentiation protocols developed by The Organoid Company. Organoids were generated and maintained in PrimeSurface® 96 Slit-well Plates (MS-9096SZ; Sumitomo Bakelite Co., Ltd., Tokyo, Japan), whose interconnected slit-well geometry permits single-step media exchange across all 96 wells without disturbing spheroid integrity, and were cultured for 90 days to achieve functional maturation prior to MPS assembly.

### Multi-organ MPS assembly

Organoids were loaded into the BioStellar™ Plate, a stirrer-pump-integrated Multi-Organ Microphysiological System (BS-X9607; Sumitomo Bakelite Co., Ltd., Japan), operated with the Stirrer Dock system (MFS-BSDOCK ver.2; Microfluidic System Works Inc., Japan). This microfluidic plate employs a kinetic pump mechanism in which integrated micro-stirrer bars, driven by the Stirrer Dock, generate continuous media perfusion through microfluidic channels connecting paired culture wells^12^. The system supports a co-culture of up to 4 organ compartments per plate. The Stirrer Dock controller permits rotation speeds of 2,500–6,000 RPM; the system was operated at the minimum setting of 2,500 RPM to minimize shear stress on organoids, as this exceeded the estimated internal hepatic metabolic processing rate and was sufficient to establish continuous media exchange between connected compartments. Three connection configurations were established: (1) Liver-CNS (main dose-response; (2) Liver-Liver (metabolite transfer control); (3) CNS-CNS (metabolism-negative control), each with n=3 biological replicates per condition.

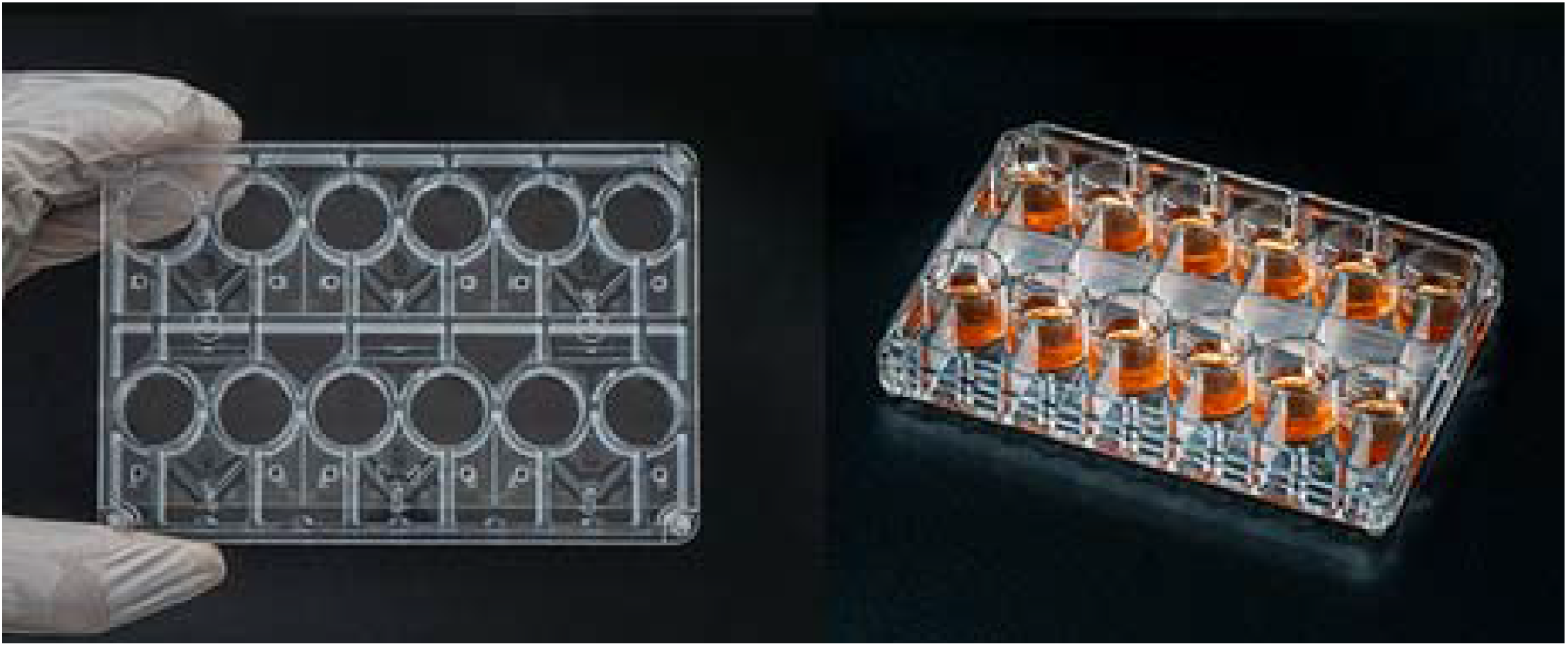
*Credit: Multi-Organ Microphysiological System BioStellar™ Plate*

### Drug treatment

Cyclophosphamide (CP) was dissolved in sterile water and applied at 0 (vehicle), 50, 100, and 200 µM to the designated treatment compartments. For Liver-CNS connections, CP was added to the liver compartment. Organoids were incubated for 48 hours under standard culture conditions (37°C, 5% CO_2_) with continuous MPS flow. Conditioned media was collected from all compartments at the 48-hour endpoint and stored at −80°C for future metabolite analysis.

### Fixation and immunostaining

After 48 hours, organoids were fixed with 4% paraformaldehyde for 30 minutes at room temperature. Immunostaining was performed using the following primary antibodies: rabbit anti-NeuN (Invitrogen, 702022; 1:500) for CNS organoids; rabbit anti-Albumin (Proteintech, 16475-1-AP; 1:500) for liver organoids; and mouse anti-γH2AX from the HCS DNA Damage Kit (Invitrogen, H10292, Component B; 1:1000) for both organoid types. γH2AX was detected with Alexa Fluor 555 goat anti-mouse secondary antibody (Kit Component C; 1:2000). Nuclear counterstaining was performed with DAPI.

### Confocal imaging

Organoids were imaged on a Yokogawa CQ1 confocal quantitative image cytometer at 10x magnification, acquiring approximately 40 Z-slices per well across 4 fields of view. Four channels were captured: brightfield, DAPI (Ch2), γH2AX/AF555 (Ch3), and NeuN or Albumin (Ch4). Fields were stitched into 2×2 mosaics and processed as volumetric Z-stacks.

### Image analysis and quantification

All image analysis was performed using custom Python scripts employing scikit-image, OpenCV, and SciPy. For each well, Z-stacks were processed slice-by-slice:

#### DAPI segmentation

Otsu thresholding was applied to each Z-slice to generate binary masks of nuclear-positive regions. The organoid core was defined as the Z-slice with maximum DAPI cross-sectional area; core metrics represent the average of the centre slice ±1 adjacent slice.

#### γH2AX quantification

A dual-metric approach was used. Puncta were detected within DAPI-masked regions using Difference-of-Gaussian (DoG) blob detection (min_sigma=1.5, max_sigma=8.0, threshold=0.015) after background subtraction. At 10x magnification (∼0.65 µm/pixel), detected puncta represent individual foci or small clusters of γH2AX foci. Mean γH2AX intensity within the DAPI mask was measured as a complementary metric capturing diffuse pan-nuclear staining. The total γH2AX signal was computed as the product of puncta density (per 1000 DAPI pixels) and mean intensity.

#### Biomarker quantification

NeuN and albumin intensity were measured within DAPI-masked regions and normalized to DAPI intensity. Biomarker-positive fraction was determined by Otsu thresholding the biomarker channel within the DAPI mask.

#### Quality control

Wells with DAPI area spanning the entire mosaic field (indicating absence of organoid and thresholding of background haze) were excluded (4 wells: A4, D6, F3, F6). Representative images for grids were selected based on quality scores incorporating organoid size, circularity, and signal-to-noise ratio, with preference for single-organoid wells.

#### Enhanced Depth-of-Field imaging

Publication images were generated using Extended Depth of Field (EDoF), which selects the best-focused pixel at each XY position across the Z-stack based on local variance of Laplacian, followed by CLAHE contrast enhancement.

### Statistical analysis

Core-level metrics were compared across dose groups using Welch’s t-test (unequal variance) relative to vehicle controls. Significance is reported as: *p<0.05, **p<0.01, ***p<0.001. Data are presented as mean ± SEM unless otherwise indicated.

## Acknowledgements

This work was supported by The Organoid Company BV, Sumitomo Bakelite Co., Ltd., and ITOCHU Corporation. The authors thank Sumitomo Bakelite Co., Ltd. for providing the BioStellar− Plate and Microfluidic System Works Inc. for the Stirrer Dock system.

## Author Contributions

T.M. and E.W. conceived the study and performed experiments, imaging, analysis, and data interpretation. All authors contributed to manuscript preparation.

